# Ribosome biogenesis is a downstream effector of the oncogenic U2AF1-S34F mutation

**DOI:** 10.1101/2019.12.13.876284

**Authors:** Abdalla Akef, Kathy McGraw, Steven D. Cappell, Daniel R. Larson

## Abstract

U2AF1 forms a heterodimeric complex with U2AF2 that is primarily responsible for 3’ splice site selection. U2AF1 mutations have been identified in most cancers but are prevalent in Myelodysplastic Syndrome and Acute Myeloid Leukemia, and the most common mutation is a missense substitution of serine-34 to phenylalanine (S34F). However, the U2AF heterodimer also has a non-canonical function as a translational regulator. Here, we report that the U2AF1 S34F mutation results in specific mis-regulation of the translation initiation and ribosome biogenesis machinery, with the potential for widespread translational changes. The net result is a global increase in mRNA translation at the single cell level. Among the translationally upregulated targets of U2AF1-S34F are Nucleophosmin1 (NPM1), which is a major driver of myeloid malignancy. Depletion of NPM1 impairs the viability of wt/S34F cells and causes rRNA processing defects, thus indicating an unanticipated synthetic interaction between U2AF1, NPM1 and ribosome biogenesis. Our results establish a unique molecular phenotype for the U2AF1 mutation which recapitulates translational mis-regulation in myeloid disease.

## Introduction

U2 Small Nuclear RNA Auxiliary Factor 1 (U2AF1; also known as U2AF35) forms a heterodimeric complex with U2AF2 (U2AF65) that is primarily responsible for 3’ splice site selection (Merendino et al., 1999). Several U2AF1 mutations have been identified in Myelodysplastic Syndrome (MDS) and Lung Adenocarcinoma (LUAD) patients (Brooks et al., 2014), the most common of which is a single missense substitution of serine-34 to phenylalanine (S34F). The mutation is heterozygous, occurs early in disease progression, and is a hotspot mutation in the RNA binding domain of the protein, strongly suggesting a gain of function, oncogenic mutation. While several groups have examined alternative splicing alterations that are mediated by the S34F mutation (Brooks et al., 2014; Fei et al., 2016; Smith et al., 2019; Yoshida et al., 2011), there is no clear set of splicing events that can explain the tumorigenic role of the S34F mutation. Previously, we reported that U2AF1, in complex with U2AF2, plays a non-canonical role in translation regulation through direct interactions with target RNA in the cytosol (Palangat et al., 2019). Recently, a U2AF1 paralog, U2AF26, has also been shown to regulate the translation of its bound mRNAs (Herdt et al., 2020). Using RNA immunoprecipitation (RIP) and polysome profiling, we found that the S34F mutation resulted in loss of binding and a concomitant increase in the translation of these target mRNAs. These mRNA targets were enriched for components of the mRNA translation and ribosome biogenesis machinery (Palangat et al., 2019). However, it is unknown how ribosome biogenesis and translational mis-regulation play a role in the physiology of U2AF1 mutations and contribute to leukemogenesis.

Intriguingly, changes in translational regulation and ribosome biogenesis are hallmarks of myeloid diseases. Hematopoietic stem cells maintain a quiescent state by limiting protein synthesis rates, and altering this balance leads to defects in cellular differentiation and lineage commitment (Signer et al., 2014). In line with this observation, germ-line mutations resulting in haploinsuffiency of ribosomal subunits such as *RPS19, RPS26, RPS17, RPL5, RPL11* or assembly chaperones such as *TSR2* (i.e. ‘ribosomopathies’) preferentially result in defects in myeloid differentiation which manifest as anemias such as Diamond-Blackfan Anemia and Shwachman-Diamond syndrome (Khajuria et al., 2018). Similarly, recent evidence suggests that MDS is characterized by increased protein synthesis in the population of CD123+ MDS stem cells from human patients (Stevens et al., 2018). These malignant MDS stem cells show increased translation yet retain a quiescent phenotype and are predominantly in the G0 phase of the cell cycle. Finally, therapeutics which target RNA polymerase I, the polymerase responsible for synthesizing rRNA, have shown specific activity against leukemia initiating cells in acute myeloid leukemia (AML) (Hein et al., 2017). In total, these studies suggest that changes in translation and ribosome biogenesis contribute to the pathology of myeloid disease and are therapeutic targets. Yet, in many cases, especially for diseases such as MDS and AML which arise from somatic mutations in stem and progenitor cells, the molecular pathways have not been described.

Among the top translationally upregulated targets of U2AF1-S34F is Nucleophosmin1 (NPM1) (Palangat et al., 2019) which plays a functional role in ribosome biogenesis and is mutated in a fifth of all AML cases. NPM1 (also known as B23) is a nucleic acid binding protein involved in the processing and nucleocytoplasmic transport of ribosomal RNA (rRNA) (Herrera et al., 1995; Savkur and Olson, 1998), (Maggi et al., 2008). Yet, despite the involvement of NPM1 in a crucial process such as ribosome biogenesis, fibroblasts derived from an *NPM1*^*-/-*^ mouse could survive in culture with modest defects (Grisendi et al., 2005). Moreover, the expression of a dominant-negative NPM1 mutant did not cause noticeable defects in the steady state levels of the 28S and 18S rRNA (Maggi et al., 2008). These findings indicate that NPM1 is not required for cell viability. The AML-linked mutations cause the mis-localization of NPM1 to the cytoplasm (Falini et al., 2005) potentially leading to an imbalance of nucleolar and cytoplasmic functions, including changes in gene expression (Brunetti et al., 2018) and p53 activity (Bertwistle et al., 2004; Itahana et al., 2003; Korgaonkar et al., 2005). Strikingly, NPM1 mutations rarely occur outside of myeloid malignancies (i.e., MDS and AML), suggesting that NPM1 plays a unique role in the differentiation and/or proliferation of myeloid cells. However, the role of NPM1 in promoting myeloid malignancies remains incompletely understood. For example, the established role that NPM1 plays in ribosome biogenesis has not been definitively linked to disease phenotypes. Moreover, despite the pervasiveness of splicing factor and NPM1 mutations in myeloid disease, there is no reported functional connection between these factors.

Here, we establish a direct link between two major drivers of myeloid disease – U2AF1 and NPM1. We find that a heterozygous U2AF1-S34F mutation leads to translational re-programming, including a global increase in protein synthesis in both immortalized epithelial cells and mouse-derived common myeloid progenitors, similar to that observed in MDS patients. Moreover, cells with the S34F mutation are dependent on NPM1 for cell-cycle progression. Depletion of NPM1 impaired the viability of the S34F-harboring cells but not wt/wt cells. Wt/S34F cells with reduced NPM1 displayed: 1) lower proliferative capacity, 2) accumulation in S phase of the cell cycle; and 3) specific rRNA processing defects. A quantitative proteomic analysis indicated that ribosome biogenesis factors and cell cycle DNA replication factors were significantly downregulated in wt/S34F cells depleted of NPM1. Our results demonstrate that NPM1 is a downstream effector of the translation changes mediated by the U2AF1-S34F mutation. Moreover, this viability defect explains why U2AF1 and NPM1 mutations are mutually exclusive in MDS and AML patients. These data indicate an unanticipated functional connection between a splicing factor mutation and the ribosome biogenesis machinery.

## Results

### The U2AF1 S34F mutation alters the level of translation initiation proteins and upregulates global translation

Using RNA immunoprecipitation followed by sequencing (RIP-seq), we previously reported that U2AF1 binds mRNA in the cytoplasm enriched for functional gene ontology (GO) categories that include “translation initiation” and “peptide biosynthetic process.” Here, we systematically address the changes in translational efficiency and total protein for those mRNA using quantitative analysis of polysome profiles and mass spectrometry. To dissect the molecular mechanisms perturbed by the U2AF1-S34F mutation, we employed three human bronchial epithelial cell lines (HBEC) (Ramirez et al., 2004) that were generated by gene-editing: (1) biallelic WT U2AF1 (herein referred to as wt/wt), (2) wt/U2AF1-S34F heterozygous (wt/S34F), and (3) a frameshift of the U2AF1-S34F allele (wt/S34F-) as previously described (Fei et al., 2016; Palangat et al., 2019).

By analyzing polysome profiling data from wt/wt, wt/S34F and wt/S34F-cells (previously published in Palangat et al., 2019), we sought to determine whether the S34F mutation altered the translational regulation of mRNA in the GO category “translation initiation” (GO:0006413). This category consists of 185 genes, overlaps many of the other categories we observed as U2AF1-bound in RIP-seq, and contains ribosomal subunits, eukaryotic initiation factors, and some ribosome biogenesis factors. Our analysis is based on a weighted estimator of translation efficiency as determined by RNA abundance in the polysome profile. Briefly, this analysis attempts to infer the number of ribosomes on an mRNA by comparing the quantitative polysome absorbance profile, the normalized counts in each fraction from RNA-seq, and the theoretical expectation for the sedimentation velocity (see methods). We applied this estimator for 20,997 RNAs which were present in each fraction of each sample in our polysome analysis (24 total measurements, Table S1). Out of 185 mRNA targets in the translation initiation GO category, 175 met these criteria, and we performed principal component analysis and hierarchical clustering on the translational efficiency computed from this estimator (Fig. 1A, B). These data indicate that the S34F mutation results in widespread changes in translation efficiency in mRNA in the GO category of translation initiation. However, these changes are almost completely rescued by frameshifting the mutant allele, as indicated by the observation that the S34F mutation accounts for > 90% of the variation in the samples (Fig. 1B). Translation efficiency both increases and decreases for genes in this family, with two distinct subgroups evident in the clustering analysis. Increases in translation efficiency occur for a number of eukaryotic initiation factors (*EIF5, EIF4E, EIF3E, EIF3H, EIF3J, EIF3M*), helicases (*DDX18, DDX1, DDX3X, DHX29*), and *NPM1*. Decreases in efficiency occur for some negative regulators (*EIF4EBP1*), members of the mTOR pathway (*RPTOR, LAMTOR2, LAMTOR4)*, and ribosomal subunits. Notably, all ribosomal subunits were either unchanged or decreased, and several genes implicated in Diamond-Blackfan Anemia (*TSR2, RPS26, RPS17*) clustered in the group showing the largest decreases in translation efficiency. Thus, a single amino acid change in the splicing factor U2AF1 results in changes in translation efficiency across messages coding for proteins which are also involved in translation initiation, with the potential for widespread changes to the proteome.

**Figure 1.**
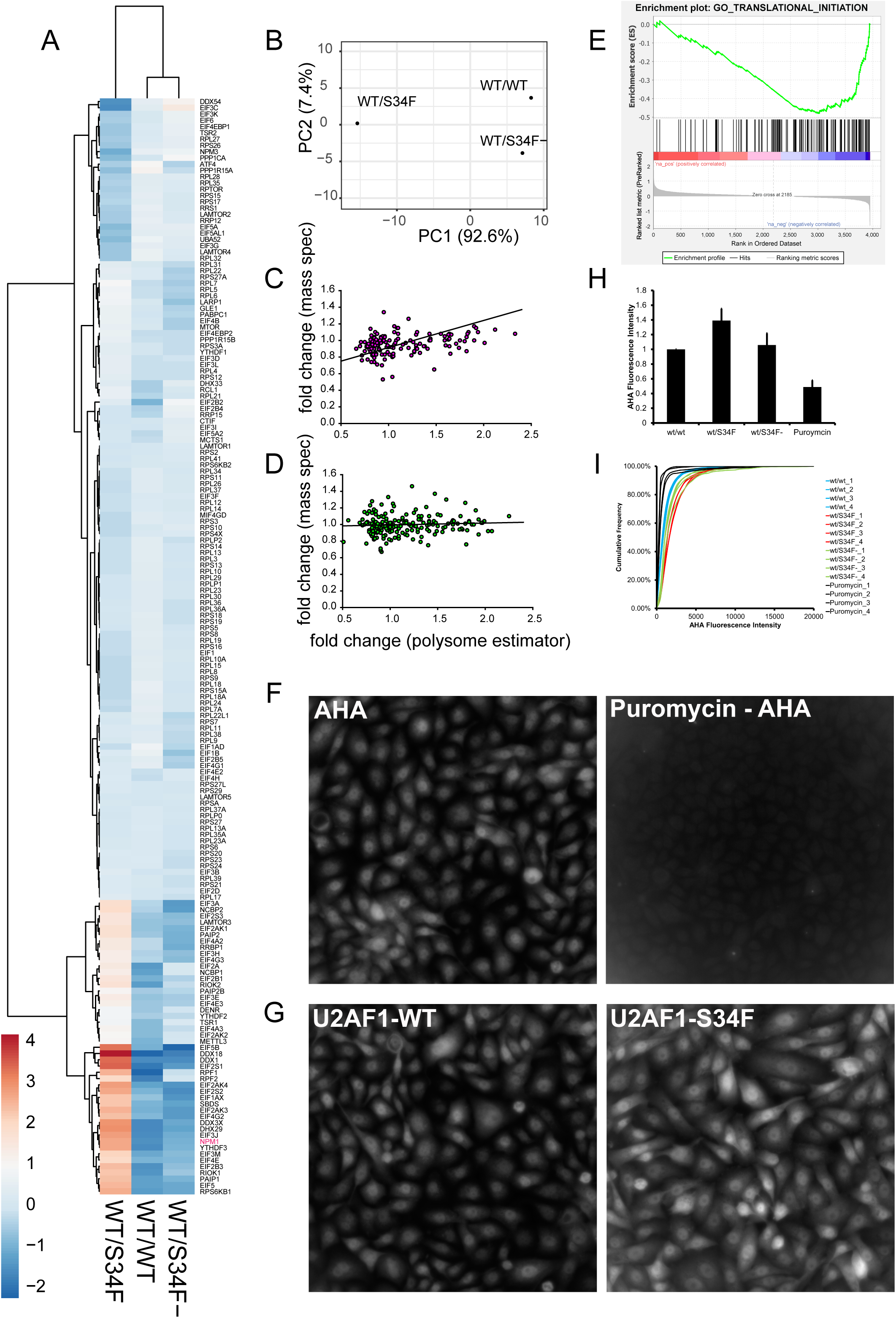
The U2AF1-S34F mutation upregulates a subset of translation and ribosome biogenesis factors and increases nascent polypeptide production. A) Heat map and hierarchical clustering of the weighted estimator applied to RNA sequencing from polysome fractions. The genes are the 175 translation initiation genes present in all eight fractions of each sample (wt/wt, wt/S34F, wt/S34F-) cells (24 total measurements). Rows are centered; no scaling is applied to rows. Rows are clustered using Euclidean distance and Ward linkage. Columns are clustered using correlation distance and average linkage. 175 rows, 3 columns. B) Principal component analysis of weighted estimator applied to RNA sequencing from polysome fractions. No scaling is applied to rows; SVD with imputation is used to calculate principal components. X and Y axis show principal component 1 and principal component 2 that explain 92.6% and 7.4% of the total variance, respectively. N = 3 data points. C) Comparison of fold change for polysome measurement to mass spectrometry measurements for genes/proteins in the “translation initiation” GO category (152 genes, Table S2). Fold change is wt/S34F over wt/wt. Regression line is reduced major axis regression with slope = 0.32 +/- 0.17, intercept = 0.59 +/- 0.04. D) Comparison of fold change for polysome measurement to mass spectrometry measurements for genes/proteins with similar abundance to proteins in panel C. Fold change is wt/S34F over wt/wt. Regression line is reduced major axis regression with slope = 0.04 +/- 0.04, intercept = 0.98 +/- 0.10. E) Gene set enrichment analysis (GSEA) based on MS data. Analysis was carried out on preranked lists using GSEA 4.0.3. Translation initiation is negatively enriched in wt/S34F cells compared to wt/wt cells. Normalized enrichment score = −2.20. FDR q-value =0.008. FWER p-value = 0.027. Full analysis in Figure S1. F) Puromycin treatment abrogates the nascent polypeptide fluorescence signal. Scale bar = 10 µm. G-H) The U2AF1-S34F mutation upregulates global nascent polypeptide production. The cells were imaged (G) and total cellular fluorescence intensity was quantified (H). Each bar represents the average and standard error of four independent experiments. p = 0.039. Scale bar = 10 µm. I) Cumulative Fluorescence intensity of the AHA signal.

To independently interrogate changes in protein levels resulting from the S34F mutation, we used quantitative mass spectrometry (MS). Wt/wt and wt/S34F cells were harvested, lysed and total protein was quantified and digested. The peptides were conjugated to tandem mass tags (TMT) and MS was performed as previously described (Rauniyar and Yates, 2014; Thompson et al., 2003). We detected 4,167 proteins present in three replicates of isobaric mass tag-MS in wt/wt and wt/S34F cells (6 total measurements). Comparing the fold change measured from MS to the fold change measured from polysome profiling across all proteins that are above the threshold in both studies (3,953 proteins, Table S2) revealed little to no correlation between the two measures. The difficulty in comparing translation efficiency as determined through sequencing approaches and MS has been reported previously and can be due to technical artifacts and also the relative contributions of protein synthesis and decay in determining total protein abundance (Ingolia, 2016). Nevertheless, for the translation initiation GO category (152 genes above threshold in polysome and MS measurements), there was a linear relationship between polysome fold-change and MS fold change (Fig. 1C, Table S2). Using reduced major axis regression, which effectively handles errors in both the *x*- and *y*-variable (Sokal and Rohlf, 2011), gives a regression coefficient of 0.32 +/- 0.17. This slope indicates that changes in translation efficiency are in fact resulting in changes in protein abundance, albeit with a smaller dynamic range. In contrast, a randomly selected group of proteins with similar abundance shows no correlation between polysome fold-change and MS fold-change (Fig. 1D, regression coefficient = 0.04).

Next, we used our MS data in wt/wt and wt/S34F as a more faithful measure of gene expression changes which arise through the S34F mutation. Since changes in translation efficiency and/or protein abundance would not be reflected in RNA abundance or isoform changes measured through RNA-seq, we performed gene set enrichment analysis on protein abundance (Subramanian et al., 2005). Positive enrichment (proteins which are increased in the wt/S34F cells compared to wt/wt cells) did not result in any gene sets with a false discovery rate (FDR) < 0.2 or family-wise error rate (FWER) < 0.2. However, analyzing the negatively enriched gene sets (proteins which are decreased in the wt/S34F cells compared to the wt/wt cells) uniquely identified genes/proteins involved in translation initiation (FDR = 0.008, FWER = 0.027, normalized enrichment score = −2.20) (Fig. 1E, Fig. S1). Thus, MS identifies changes in translation initiation proteins as the dominant molecular phenotype for the U2AF1 S34F mutation.

Finally, we sought to examine the functional consequences of mis-regulation of the translation initiation machinery at the single-cell level. Our genome-wide assays pointed in multiple directions: translation initiation as a category was negatively enriched in wt/S34F cells compared to wt/wt cells, but this category encompasses both positive and negative regulators of initiation. To measure the integrated output of these changes, we compared global mRNA translation rates between the wt/wt and the wt/S34F cells using the fluorescent noncanonical amino acid tagging (FUNCAT) as previously described (Tom Dieck et al., 2012). Cells were pulsed with the methionine analog azido-homoalanine (AHA) for 1 hour. Subsequently, cells were fixed, permeabilized and the AHA-containing nascent polypeptides were labelled for microscopy using click chemistry. First, we confirmed that puromycin treatment abolished the AHA fluorescence signal (Fig. 1F). Next, we found that the S34F mutation caused an upregulation in global translation (Fig. 1G, quantification in Fig.1H). Comparing fluorescence intensity at the single cell level indicated that the elevated level of global translation in the wt/S34F cells was representative of the whole cell population rather than a few outlier cells (Fig. 1I). Taken together, we conclude that the mis-regulation of the translation initiation machinery in wt/S34F cells results in an increase in global mRNA translation.

In summary, a single missense mutation in U2AF1 results in widespread changes to translation as measured both by polysome sequencing and mass spectrometry. Proteins involved in translation initiation -- including eukaryotic initiation factors, ribosome subunits, positive and negative translational regulators, and ribosome biogenesis factors – are significantly affected. Despite a decrease in protein abundance for the GO category of translation initiation as a whole in wt/S34F cells compared to wt/w/t cells, protein synthesis is elevated in these cells, recapitulating the phenotype seen in human MDS stem cells (Stevens et al., 2018).

### Wt/S34F cells require NPM1 for survival in a p53-independent manner

Because NPM1 was identified first as a target of U2AF1 binding in the cytosol, is included in the translation initiation gene set, shows concordant increase in translational efficiency by polysome sequencing and protein abundance by MS (Table S2), and is frequently altered in myeloid malignancy, we chose to focus on this mRNA/protein for in depth characterization. We first confirmed the upregulation of NPM1 translation in wt/S34F cells using a dual luciferase reporter. The first exon of *NPM1* was fused upstream to Renilla luciferase. To normalize for differences in cell number and copy number, the reporter also included a Firefly luciferase downstream of an Internal Ribosome Entry Site (IRES) sequence derived from the Cricket Paralysis Virus (CrPV) (Coller and Parker, 2005; Fernández et al., 2014) (Fig. 2A). The wt/S34F cells had an 8-fold higher Renilla to Firefly luciferase activity than wt/wt or wt/S34F-cells (Fig. 2B) further confirming the upregulation of NPM1 in cells harboring the U2AF1-S34F mutation and additionally indicating that the 5’ end of the mRNA is sufficient for increased translation.

**Figure 2.**
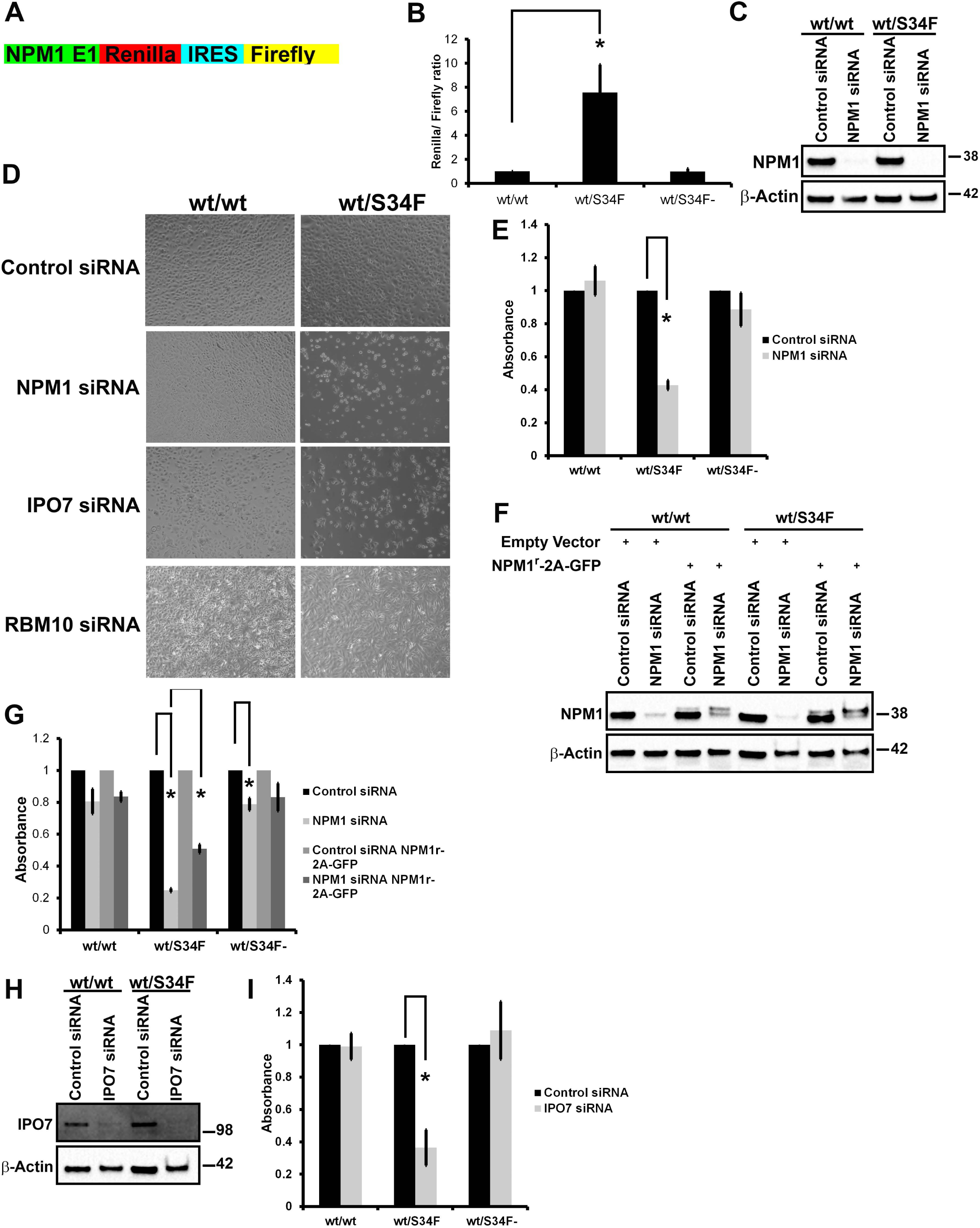
NPM1 and IPO7 are required for the viability of the wt/S34F cells. A) A schematic diagram of the *NPM1-luciferase* reporter. The reporter consists of NPM1 exon 1, Renilla luciferase, CrPV IRES, Firefly luciferase. B) Normalized Renilla to Firefly luciferase activity demonstrates elevated translation levels of the *NPM1-luciferase* reporter. p = 0.009. C) Cells treated with siRNA directed against NPM1 were lysed and proteins separated on SDS-PAGE and analyzed by immunoblot using antibodies against NPM1 and α-Tubulin. D) Cells treated with the indicated siRNAs were imaged 144 hours post-siRNA transfection. Scale bar = 50 µm. E) Quantification of cell viability using the WST1 reagent. Each bar represents the average and standard error of three independent experiments. p = 0.019 using a paired 2-sided t-test. F) Cells that express either NPM1^r^-2A-GFP or empty vector were treated with siRNA directed against NPM1 were lysed and proteins separated on SDS-PAGE and analyzed by immunoblot using antibodies against NPM1 and β-Actin. G) Quantification of cell viability using the WST1 reagent. Each bar represents the average and standard error of three independent experiments. * = p < 0.05 using a paired 2-sided t-test. H) Cells treated with siRNA directed against IPO7 were lysed and proteins separated on SDS-PAGE and analyzed by immunoblot using antibodies against IPO7 and β-Actin. I) Quantification of cell viability using the WST1 reagent. Each bar represents the average and standard error of three independent experiments. p = 0.028 using a paired 2-sided t-test.

We sought to examine whether increased NPM1 translation was a downstream mediator of the pro-tumorigenic phenotype of the U2AF1-S34F mutation. NPM1 was depleted by RNAi in wt/wt, wt/S34F and wt/S34F-cells (Fig 2C). We noticed a marked decrease in the viability of the wt/S34F cells upon NPM1 depletion (Fig. 2D) concomitant with a decrease in metabolic activity (Fig 2E). No defects were observed for the wt/wt or wt/S34F-cells (Fig. 2D, E). We generated cell lines that express an RNAi-resistant NPM1 construct (NPM1^r^-2A-GFP) using lentiviral-mediated integration. A slower mobility band was detected for NPM1^r^-2A-GFP that was unaffected by NPM1 siRNA treatment (Fig. 2F). The viability of the wt/S34F cells that express NPM1^r^-2A-GFP was partially restored (Fig. 2G). The partial rescue is likely due to the low expression levels of NPM1^r^-2A-GFP in comparison to endogenous NPM1 levels (Fig. 2F).

We sought to corroborate these findings by depletion of a second ribosome biogenesis factor, Importin 7 (IPO7). IPO7 mRNA is also a direct U2AF1 target in the cytoplasm and is responsible for the import of the large ribosomal subunit proteins RPL5 and RPL11 to the nucleus (Golomb et al., 2012). As for NPM1, IPO7 silencing by RNA interference (RNAi) was also not reported to cause cell death (Golomb et al., 2012). Yet, IPO7 depletion also specifically impaired the survival of wt/S34F but not wt/wt cells (Fig. 2D, H-I). Neither a scrambled control siRNA nor siRNA to RBM10, which is a splicing factor in the nucleus, had any effect on cell viability in either cell line (Fig. 2D). Taken together, NPM1 and IPO7, both targets of U2AF1 binding and translational regulation (Table S1), both involved in transport and/or ribosome biogenesis, are essential for viability in the wt/S34F cells but not in the wt/wt cells.

It is known that disruption of ribosome biogenesis and/or alteration of the stoichiometry of ribosome subunits can trigger a p53 response in cells (Ajore et al., 2017; Dutt et al., 2011). Specifically, NPM1 downregulates the activity of p53 by sequestering ARF (also known as p14) in the nucleolus and preventing the ARF-MDM2 interaction, thus enabling MDM2-mediated degradation of p53 (den Besten et al., 2005; Korgaonkar et al., 2005; Mariano et al., 2006). We therefore investigated whether the viability defect of the wt/S34F cells depleted of NPM1 was dependent on p53 activity. First, we compared p53 levels between wt/wt and wt/S34F cells depleted of NPM1. NPM1 depletion caused the upregulation of p53 in both the wt/wt and wt/S34F cells to a comparable extent (Fig. 3A). Moreover, the synergistic depletion of NPM1 and p53 did not rescue the viability defect of the wt/S34F cells caused by the absence of NPM1 (Fig 3B, C). Next, we sought to assess if increasing p53 levels in an NPM1-independent mechanism would specifically impair the viability of the wt/S34F cells. p53 levels were induced by treating cells with 10µM Nutlin3A (Tovar et al., 2006) (Fig. 3D). p53 activation by increasing Nutlin3A concentration impaired the viability of wt/wt, wt/S34F and wt/S34F-cells to a similar extent (Fig. 3E, quantification in Fig. 3F). Thus, we conclude that the depletion of NPM1 impairs the viability of the wt/S34F cells through a p53-independent pathway.

**Figure 3.**
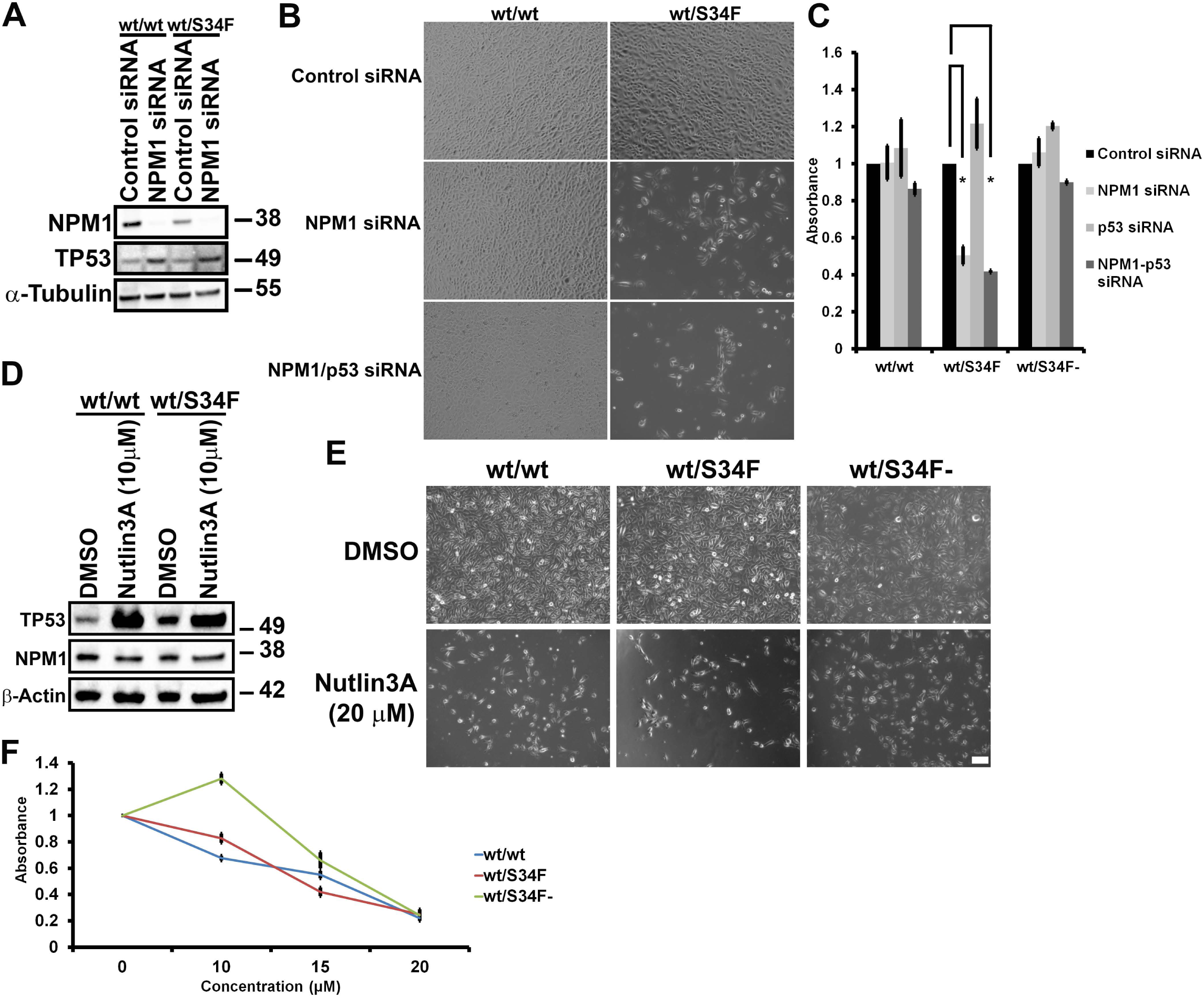
The requirement of wt/S34F cells for NPM1 is independent of TP53. A) Cells treated with siRNA directed against NPM1 were lysed and proteins separated on SDS-PAGE and analyzed by immunoblot using antibodies against NPM1 and α-Tubulin. B) Cells treated with the indicated siRNAs were imaged 144 hours post-siRNA transfection. Scale bar = 50 µm. C) Quantification of cell viability using the WST1 reagent. Each bar represents the average and standard error of three independent experiments. * = p < 0.05 using a paired 2-sided t-test. D) Cells treated with 10µM Nutlin3A for 36 hours were lysed and proteins separated on SDS-PAGE and analyzed by immunoblot using antibodies against NPM1, p53 and β-Actin. E) Nutlin3A treatment impairs the viability of wt/wt, wt/S34F and wt/S34F-cells. Scale bar = 50 µm. F) Cell viability was assessed 36 hours after treatment with the indicated doses of Nutlin3A using the WST1 reagent.

### Wt/S34F cells depleted of NPM1 exhibit cycle progression defects and changes in ribosome biogenesis factors

The viability defect observed in wt/S34F cells lacking NPM1 can be explained by either an activation of apoptotic pathways or a defect in cell cycle progression. To distinguish between these possibilities, we first examined cellular viability over the time course of the knock-down experiment. First, wt/wt cells transfected with control siRNA had comparable growth with wt/wt cells depleted of either NPM1 or IPO7. In contrast, wt/S34F cells depleted of NPM1 or IPO7 had a drastically lower viability than wt/S34F cells treated with control siRNA (Fig. 4A), consistent with our earlier observations (Fig. 2D). Importantly, we observed a monotonic increase in the metabolic activity of the wt/S34F cells depleted of NPM1 or IPO7 over the time period of the measurement which is more consistent with slowed growth rates and not an induction of apoptotic pathways. We further confirmed the lack of apoptosis by comparing Active Caspase-3 levels. We chose Caspase-3 since it is one of the late apoptotic factors and thus is activated by both the extrinsic and intrinsic apoptotic pathways. We did not observe a difference in active Caspase-3 levels between wt/wt and wt/S34F cells treated with either control or NPM1 siRNA suggesting that apoptosis is not the cause of decreased viability in the wt/S34F cells lacking NPM1 activity (Fig. 4B, Supplementary Fig. 2A). In contrast, ActinomycinD treatment (5ug/mL) caused a robust upregulation in Caspase-3 levels.

**Figure 4.**
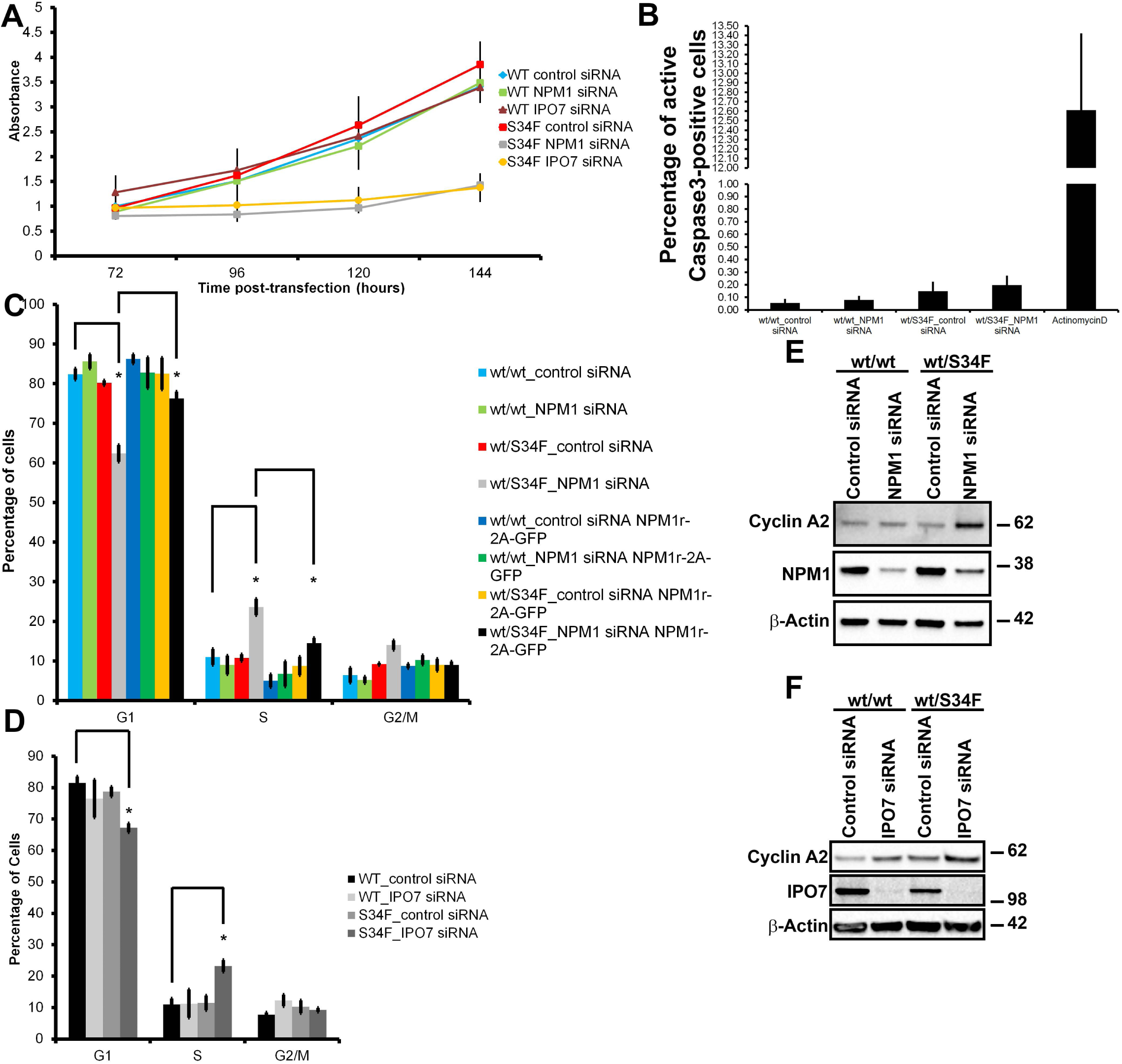
NPM1 or IPO7 silencing slows the proliferation of S34F cells. A) The viability of cells treated with siRNAs directed against NPM1 or IPO7 was assessed at the indicated timepoints post-siRNA transfection. Error bars represent the standard error of three independent experiments. B) Quantification of the active Caspase-3 levels. Each bar represents the average and standard error of three independent experiments. C) Cells that express either NPM1^r^-2A-GFP or empty vector were treated with siRNA directed against NPM1. 144 hours post-siRNA transfection, cells were pulsed with EdU for 10 minutes and EdU incorporation was quantified. The fraction of cells in each phase of the cell cycle was calculated. Each bar represents the average and standard error of three independent experiments. * = p < 0.05 using a paired 2-sided t-test. D) Cells were treated with siRNA directed against IPO7. 144 hours post-siRNA transfection, cells were pulsed with EdU for 10 minutes and EdU incorporation was quantified. The fraction of cells in each phase of the cell cycle was calculated. Each bar represents the average and standard error of three independent experiments. p = 0.002 using a paired 2-sided t-test. E-F) Cells treated with siRNA directed against NPM1 (E) or IPO7 (F) were lysed and proteins separated on SDS-PAGE and analyzed by immunoblot using antibodies against Cyclin A2, NPM1 or IPO7 and β-Actin.

Next, we examined whether wt/S34F cells depleted of NPM1 had cell cycle progression defects. Cells were treated with 5-ethynyl-2’-deoxyuridine (EdU) for 10 minutes to label the S phase population (Salic and Mitchison, 2008). The G1 and G2 cell populations were identified by quantifying the DNA fluorescence intensity using Hoechst (Fig. 4C, Supplementary Fig. 2B). The depletion of NPM1 in wt/wt cells did not have a measurable effect on cell cycle progression. In contrast, the absence of NPM1 from the wt/S34F cells led to an increase in the fraction of S phase cells and a concomitant decrease in the G1 population (Fig. 4C). These results indicate that wt/S34F cells depleted of NPM1 take a longer time to complete S phase. Expression of NPM1^r^-2A-GFP in wt/S34F when NPM1 is silenced restored the S phase population to control siRNA levels (Fig. 4C). We also observed elevated levels of the S phase population in wt/S34F cells when IPO7 was silenced (Fig. 4D, Supplementary Fig. 2C). We further confirmed the accumulation of cells in S phase by examining the levels of Cyclin A2 (CCNA2), which is upregulated in S, G2 and early M phase. Indeed, Cyclin A2 levels were specifically upregulated in wt/S34F cells depleted of NPM1 (Fig. 4E) or IPO7 (Fig. 4F). Silencing NPM1 or IPO7 expression in wt/wt cells did not have a detectable effect on Cyclin A2 levels. In summary, the absence of NPM1 or IPO7 caused the accumulation of wt/S34F cells in S phase leading to slower growth and a proliferative disadvantage.

In order to dissect the molecular pathways that are perturbed in wt/S34F cells depleted of NPM1, we repeated our quantitative MS assays after NPM1 knockdown. Of a total of 4,380 proteins detected in three replicates across two samples (6 total measurements), NPM1 depletion in wt/wt cells had a minimal effect on the cellular proteome. Only 7 and 13 proteins were up- and down-regulated respectively above a fold change of 1.5. Importantly, these factors were not significantly enriched in specific GO categories. In contrast, NPM1 silencing in wt/S34F caused 26 and 152 proteins to be up- and down-regulated respectively. A GO term analysis indicated that the downregulated factors were statistically enriched for “DNA replication coupled to cell cycle” and “ribosome biogenesis” (Fig. 5A). The downregulation of factors that couple DNA replication to cell cycle progression corroborates the accumulation of wt/S34F cells depleted of NPM1 in the S phase (Fig. 4C, D). The negative enrichment for ribosome biogenesis factors suggested a direct mechanistic link to NPM1 function and the translational mis-regulation which arises from the S34F mutation, spurring us to investigate this observation in greater detail.

**Figure 5.**
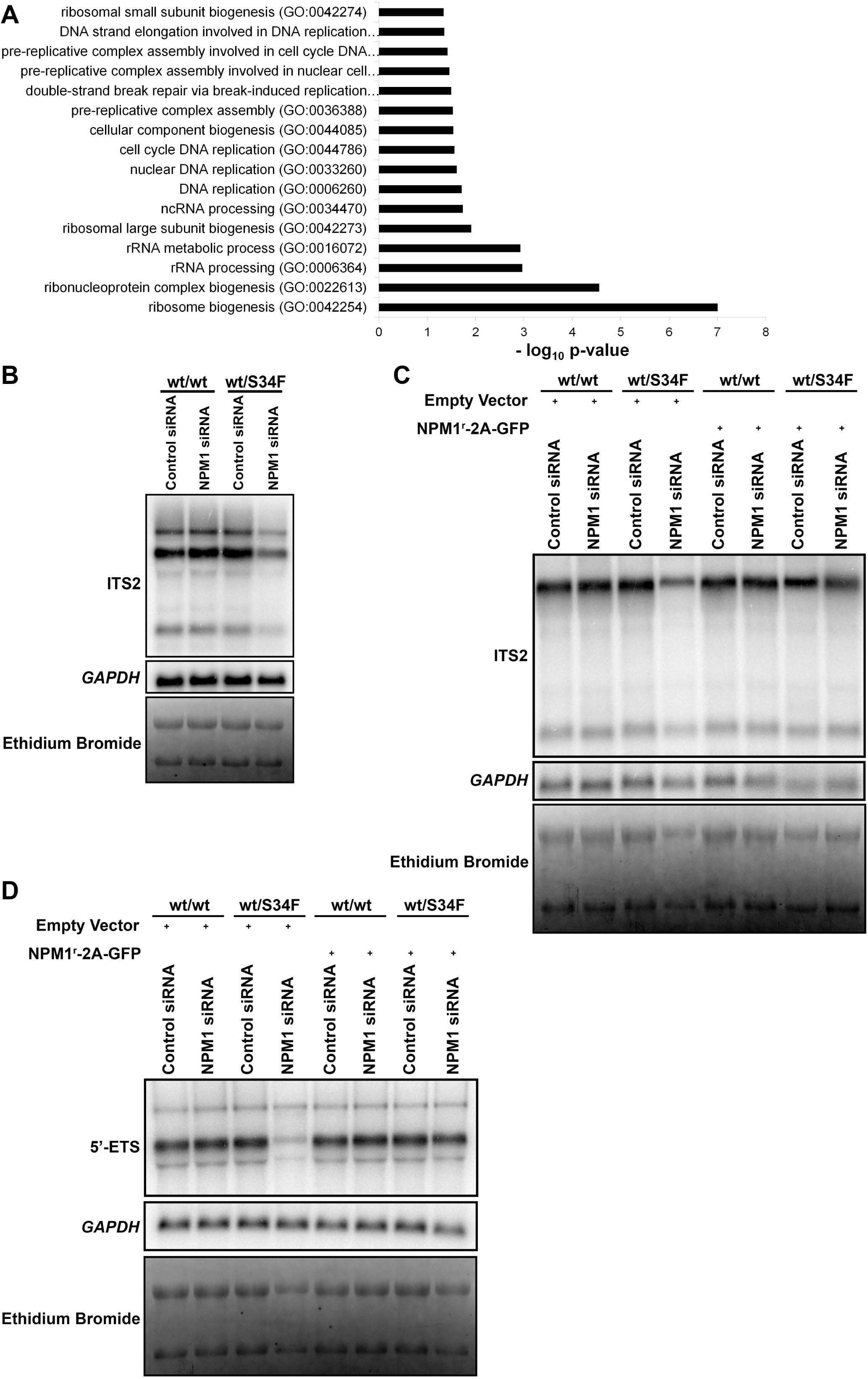
Silencing NPM1 or IPO7 impairs rRNA precursor levels in wt/S34F cells. A) Statistically enriched GO terms for downregulated proteins in wt/S34F cells depleted of NPM1 in comparison to wt/wt cells. B) The levels of the ITS2-containing rRNA precursors are decreased in wt/S34F cells depleted of NPM1. C) Overexpression of the RNAi-resistant NPM1^r^-2A-GFP restores the levels of the ITS2-containing rRNA precursors. D) NPM1 silencing in wt/S34F decreases the level of the 47S rRNA and is rescued upon expression of the RNAi-resistant NPM1^r^-2A-GFP construct.

### Processing of the 28S rRNA precursors is impaired in wt/S34F cells depleted of NPM1

Ribosome biogenesis is initiated when RNA polymerase I transcribes ribosomal DNA (rDNA) into the 47S pre-ribosomal RNA (rRNA) transcript that encompasses the 18S, 5.8S and the 28S flanked by 2 external (5’ external transcribed spacer (5’ ETS) and 3’ ETS) and interspersed by 2 internal spacers (internal transcribed spacer 1 (ITS1) and ITS2). The 18S rRNA and small subunit ribosomal proteins constitute the pre-40S ribosomal subunit. The 28S, 5.8S, and 5S (produced by RNA polymerase III) along with the large subunit ribosomal proteins constitute the pre-60S subunit. NPM1 has been previously shown to bind the 28S rRNA (Huang et al., 2005) and regulate the processing of the 28S rRNA by promoting the localization of the ITS2-cleaving endoribonuclease LAS1L to the nucleolus (Castle et al., 2012). Therefore, we reasoned that silencing NPM1 might cause a specific impairment in the processing of the 28S rRNA precursors.

We examined the levels of the 28S rRNA precursors by employing Northern blot using a probe complementary in sequence to the ITS2 (Donati et al., 2013). Silencing NPM1 in wt/wt cells did not cause a change in the levels of the ITS2-containing precursors. In contrast, the levels of the ITS2 precursors in wt/S34F cells decreased after NPM1 was depleted (Fig. 5A). The levels of *GAPDH* mRNA demonstrate equal loading across samples. However, the decrease in the ITS2 precursors did not cause a noticeable change in mature 28S and 18S rRNA levels likely due to their long half-life ranging from three to seven days (Gillery et al., 1995; Halle et al., 1997). Furthermore, the decrease in ITS2 levels in wt/S34F cells in the absence of NPM1 is rescued by expressing the RNAi-resistant NPM1^r^-2A-GFP (Fig. 5B).

The decrease in ITS2 levels can be explained by two possibilities: 1) a downregulation in rRNA transcription upon silencing NPM1 in wt/S34F cells; or 2) the ITS2-containing rRNA precursors are more rapidly processed into the mature 28S rRNA. To identify the mechanism underlying the decreased levels of the ITS2-containing precursors, we examined the levels of the primary 47S rRNA using a probe that is complementary to the 5’ ETS (Fumagalli et al., 2009). Since the 5’ ETS is present in the 47S but is rapidly cleaved during the early processing stages, the 5’ ETS containing rRNA precursors provide a direct measure of rDNA transcription. Silencing NPM1 in wt/S34F cells caused a marked decrease in the 47S rRNA (Fig. 5C). In contrast, 47S rRNA was comparable between wt/wt cells, wt/S34F cells and wt/wt cells depleted of NPM1. These results indicate that rDNA transcription is impaired in wt/S34F cells depleted of NPM1 leading to a decrease in the levels of the 47S (Fig. 5C) and 32S (Fig. 5A-B) rRNA precursors. Thus, wt/S34F cells appear to rely on NPM1 for sufficient rDNA transcription.

### The U2AF1 S34F mutation upregulates global translation in mouse myeloid progenitors and is mutually exclusive with NPM1 mutations in human patients

Our previous studies were carried out in human bronchial epithelial cells. These cells have the benefit of being immortalized non-transformed human cell lines which might better phenocopy the context in which initiating mutations such as U2AF1 S34F occur. However, U2AF1 mutations (like all splicing factor mutations) occur most frequently in the context of myeloid malignancy, so we examined the functional relevance of these findings with two additional approaches: experimental perturbation of immortalized mouse myeloid progenitor cells in tissue culture and analysis of human patient somatic mutation data.

First, we sought to examine whether the U2AF1-S34F mutation upregulates mRNA translation in myeloid cells. We employed Hoxb8 overexpression to immortalize myeloid progenitor cells as previously described (Wang et al., 2006). These myeloid progenitors were derived from bone marrow of transgenic mice that express the human cDNA of either U2AF1-WT or U2AF1-S34F from the Col1a1 locus (Shirai et al., 2015). Hoxb8 is expressed as a fusion to the estrogen-binding domain of the estrogen receptor (ER). In the presence of estrogen, Hoxb8 arrests myeloid differentiation and maintain the cells at the granulocyte/macrophage progenitor (GMP) stage. Upon estrogen withdrawal, the cells differentiate into neutrophils and macrophages that express the pan-myeloid marker CD11b (Wang et al., 2006). Strikingly, we observed that the U2AF1-S34F mutation impairs myeloid differentiation and maintains the mutant cells in a less differentiated state. In the absence of estrogen, i.e. under differentiation conditions, about 70% of wt/wt cells expressed the pan-myeloid differentiation marker CD11b. In contrast, only 45% of wt/S34F cells expressed CD11b (Fig. 6A-B). Thus, wt/S34F GMP cells are not able differentiate to the same extent as wt/wt cells, indicating that the S34F missense mutation is sufficient to recapitulate aspects of the MDS cytopenia phenotype *in vitro*.

**Figure 6.**
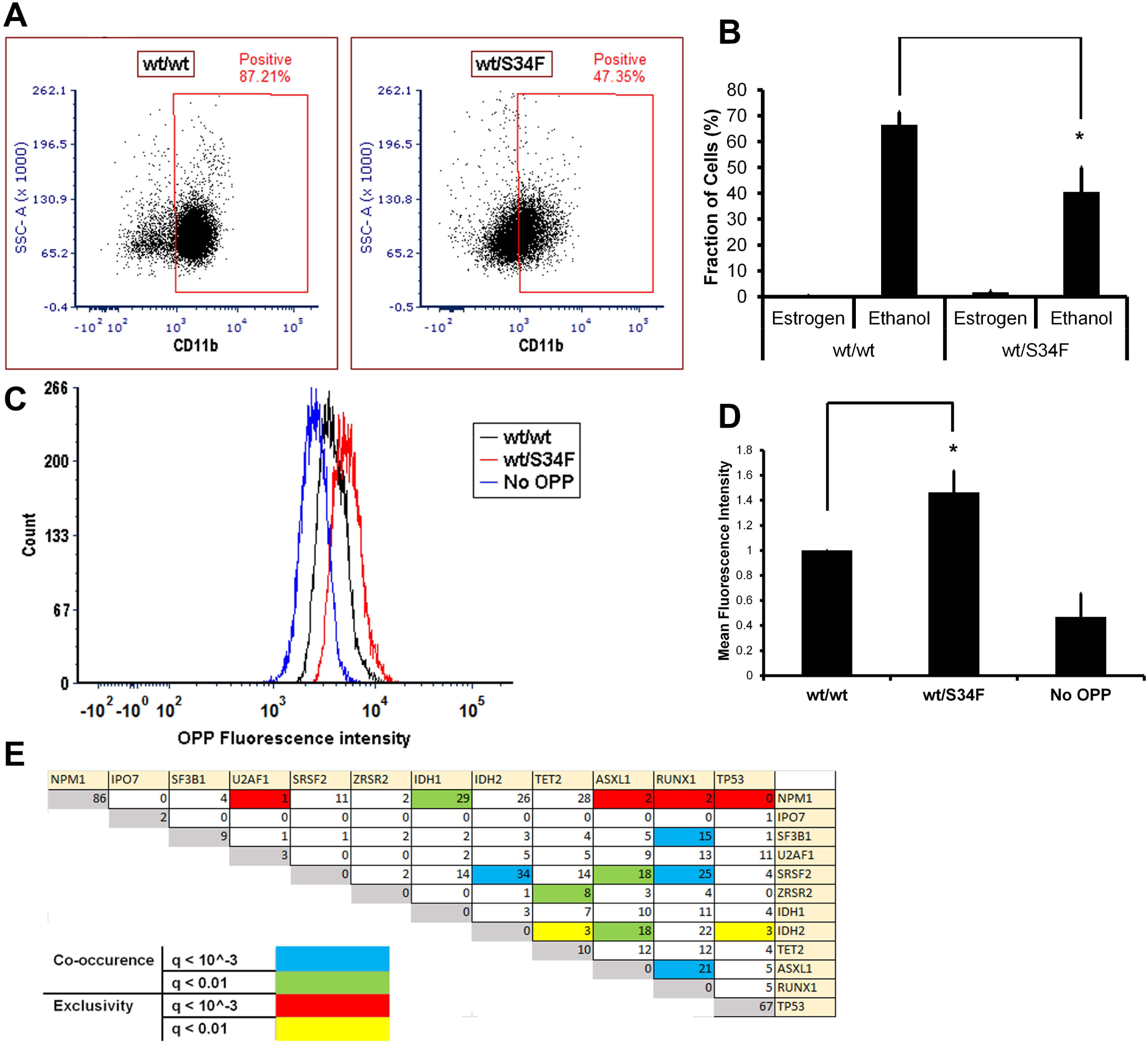
The U2AF1-S34F mutation upregulates global translation in immortalized myeloid cells. A) The U2AF1-S34F mutation impairs the ability of myeloid progenitor cells to differentiate. B) Quantification of A. Each bar represents the average and standard error of three independent experiments. p = 0.042. C) The U2AF1-S34F mutation upregulates global mRNA translation. D) Quantification of C. Each bar represents the average and standard error of four independent experiments. * = p < 0.05. E) U2AF1 and NPM1 mutations are mutually exclusive in MDS and AML patients (q value < 0.001).

Next, we examined global mRNA translation in wt/wt and wt/S34F myeloid progenitor cells. We treated the myeloid progenitors with the puromycin analog, O-propargyl-puromycin (OPP), which incorporates into nascent polypeptide chains. The OPP-containing polypeptides are then chemically reacted to a fluorophore and detected by flow cytometry as previously described (Signer et al., 2014). This assay is functionally similar to the AHA assay used previously but works better for suspension cells. Similar to HBECs, wt/S34F myeloid progenitors had higher global mRNA translation in comparison to wt/wt cells (Fig. 6C-D). Again, the elevated global mRNA translation levels in wt/S34F cells shows that the U2AF1 S34F mutation in progenitor cells *in vitro* is sufficient to recapitulate the elevated global translation observed in hematopoietic stem cells derived from MDS patients (Stevens et al., 2018).

Second, since the S34F cells require NPM1 for their viability, we hypothesized that the S34F mutation should not co-occur in cancer patients with NPM1 loss of function mutations. Both *U2AF1* and *NPM1* are frequently mutated in patients with malignancies of the myeloid lineage. According to the cBioPortal database, *U2AF1* mutations occur in 5% and *NPM1* is mutated in 22% of all myeloid malignancy patients. Despite the high frequency of these mutations, we found that *U2AF1* and *NPM1* mutations co-occurred at a probability lower than that expected by chance (Fig. 6E), likely due to the deleterious viability defect caused by abolishing NPM1 function in cells harboring U2AF1 mutations. This analysis recapitulated other previously observed mutual dependencies. For example, *SRSF2* and *IDH2* are more likely to co-occur (Yoshimi et al., 2019). Notably, *NPM1* mutations were mutually exclusive with *U2AF1* mutations but not with other commonly mutated splicing factors such as *SF3B1, SRSF2* and *ZRSR2*. Taken together, these analyses indicate that both the role of U2AF1 S34F in translation regulation and the interaction between U2AF1 and NPM1 are relevant in a disease context.

## Discussion

Although U2AF1 mutations have been implicated as driver mutations in lung cancer and acute myeloid leukemia, the role of these mutations in disease progression has remained elusive. Here, we demonstrate an unanticipated functional connection between U2AF1, translation initiation, and ribosome biogenesis. We show that the U2AF1-S34F mutation leads to a substantial change in the translation machinery resulting in an increase in global translation at the single cell level in both human bronchial epithelial cells and mouse myeloid progenitor cells, thus defining a unique molecular phenotype for this splicing factor mutation. Furthermore, wt/S34F cells require ribosome biogenesis factors such as NPM1 and IPO7 to proliferate. A quantitative proteomic analysis demonstrates that silencing NPM1 in wt/S34F cells causes a statistically significant decrease in factors that regulate ribosome biogenesis and cell cycle progression, and this observation is corroborated by a concomitant decrease in rRNA production. This *in vitro* ‘non-oncogene addiction’ of the S34F mutation to NPM1 is validated by human patient data demonstrating that *U2AF1* and *NPM1* mutations are mutually exclusive in myeloid malignancies.

To our knowledge, this U2AF1 S34F mutation is the first missense mutation to recapitulate a translation phenotype in myeloid malignancy. Although the translation machinery has long been known to be misregulated in cancer (Ruggero and Pandolfi, 2003) and is the molecular basis for the inherited ribosomeopathies which affect myeloid differentiation, these latter genetic alterations are typically germ-line deletions of ribosomal subunits. Somatic mutations in the translation machinery or related translation initiation factors have not been reported in MDS or AML. Importantly, we show this phenotype in the context of immortalized mouse myeloid progenitors in which the S34F mutation is able to phenocopy MDS-linked symptoms such as altered differentiation leading to cytopenias, anemias and neutropenias. However, the molecular changes in the abundance of translation initiation proteins induced by the S34F mutation are not easily interpreted. For example, this family of proteins is decreased in comparison to wt/wt cells, but not evenly so. A number of eukaryotic initiation factors, helicases and NPM1 are elevated, while ribosome subunits are generally down. Yet, the net output is increased peptide biosynthesis as measured by the incorporation of the amino acid analog (AHA) or tRNA analog (OPP). The change to translational activity is evocative of numerous studies showing that translational regulation is particularly important during hematopoiesis (Khajuria et al., 2018; Signer et al., 2014; Stevens et al., 2018; van Galen et al., 2018). The nature of this increased translation that we and others observe is still unclear. Are certain functional categories up-regulated? Does the increased translation result in full length or truncated proteins? Additional work is necessary to precisely answer these questions.

Previous work has identified p53 as an important mediator of ribosome dysfunction, especially in bone-marrow failure (Dutt et al., 2011; Fumagalli et al., 2012). For example, p53 accumulates in the erythroid lineage upon knockdown of RPS14, the ribosomal protein gene deleted in a subtype of MDS called 5q-syndrome (Dutt et al., 2011). However, erythroid differentiation defects could be rescued by pharmacological inhibition of p53. However, while NPM1 has been reported to regulate p53 activity either directly (Colombo et al., 2002) or through ARF (Bertwistle et al., 2004; Itahana et al., 2003; Korgaonkar et al., 2005), several lines of evidence argue against a role for p53 in the viability defect of S34F cells depleted of NPM1. First, abrogating NPM1 expression causes comparable levels of p53 induction in both the wt/wt and wt/S34F cells (Fig. 3A). Second, co-depletion of NPM1 and p53 did not rescue the viability defect observed in single NPM1 depletion (Fig. 3B, C). Third, p53 activation by 20uM Nutlin3A caused a comparable impairment of the viability of wt/wt and wt/S34F cells (Fig. 3D-F). Thus, we conclude that the phenotype we observe, namely the dependence of the S34F mutation on functional NPM1, is p53-independent.

Why are ribosome biogenesis factors specifically required for the viability and proliferation of wt/S34F cells but not wt/wt U2AF1 cells? We note that these experiments were performed in human bronchial epithelial cells, which are an immortalized but non-transformed cell line (Ramirez et al., 2004). This system might better recapitulate splicing factor mutations, which usually occur early in disease progression (Abelson et al., 2018). In tissues, cells must strike a balance between proliferation, differentiation, and self-renewal, for example as demonstrated in hematopoiesis (Khajuria et al., 2018). This balance is altered in MDS, and hematopoietic stem cells derived from MDS patients have elevated mRNA translation levels (Stevens et al., 2018). Our isogenic cell culture system recapitulates this finding with only a single nucleotide change in the *U2AF1* gene. Moreover, we speculate that ribosome biogenesis factors such as NPM1 or IPO7 which are not required for cell viability under basal growth levels (Golomb et al., 2012; Grisendi et al., 2005) become crucial when mRNA translation levels are elevated due to the U2AF1-S34F mutation. The absence of NPM1 impairs the production of the 47S pre-rRNA precursor in wt/S34F cells and this slows cell proliferation and causes their accumulation in S phase. Indeed, NPM1 has been shown to localize to centrosomes and regulate their duplication suggesting potential links between ribosome biogenesis and cell division factors (Grisendi et al., 2005; Okuda et al., 2000). Moreover, mTOR, another growth-promoting signaling pathway, has been shown to upregulate NPM1 expression (Pelletier et al., 2007). It should be also noted that abrogating the expression of the small ribosome subunit protein, RPS6 impairs cell division in mice (Volarevic et al., 2000).

Our results suggest a model where the U2AF1-S34F mutation alters the translation of mRNA that code for many translation and ribosome biogenesis leading to a global increase in protein synthesis (Fig. 1). While NPM1 silencing was not sufficient to perturb cell viability (Grisendi et al., 2005; Maggi et al., 2008) or rRNA processing (Maggi et al., 2008) in various cell lines that express WT U2AF1, NPM1 silencing has been shown to alter nucleolar shape (Nicolas et al., 2016) and the localization of ribosome biogenesis factors to the nucleolus (Castle et al., 2012). We propose increased cellular dependence on NPM1 under conditions of elevated mRNA translation such as in the presence of U2AF1-S34F. NPM1 silencing in wt/S34F cells impairs rRNA transcription (Fig. 5B-D) and ribosome biogenesis factors are downregulated (Fig. 5A) potentially at the transcriptional, translational or post-translational levels in wt/S34F cells. Furthermore, the downregulation of factors that couple DNA replication to cell cycle progression causes the accumulation of wt/S34F cells depleted of NPM1 in S phase (Fig. 4) and their decreased viability (Fig. 2). The dependence of wt/S34F cells on NPM1 accounts for the mutual exclusivity of U2AF1 and NPM1 mutations in MDS and AML patients (Fig. 6E). In sum, this study uncovered a novel dependence of the cancer-associated U2AF1 mutations on ribosome biogenesis to maintain a highly proliferative state.

## Materials and Methods

### Plasmids and cells lines

The plasmid expressing the NPM1^r^-2A-GFP reporter was generated by fusing the cDNA of NPM1 amplified from HBEC cDNA library upstream of 2A-GFP synthetic construct (IDT). This was subsequently inserted into a lentiviral transfer plasmid that has the Ubiquitin C promoter. The NPM1 luciferase reporter was generated by fusing 2 synthetic constructs (IDT) into the Ubiquitin C lentiviral transfer plasmid. The first construct was composed of human NPM1 exon 1 sequence fused to Renilla luciferase. The second construct composed of CrPV IRES fused to firefly luciferase (Coller and Parker, 2005). The lentiviral plasmids were packaged into lentiviral particles using HEK293T cells. Viruses were collected from the supernatant and transduced into HBEC using 8ug/mL polybrene. HBEC lines were cultured in keratinocyte serum-free medium supplemented with bovine pituitary extract and epidermal growth factor (Invitrogen). Wt/wt, wt/S34F and wt/S34F-HBEC cells were previously reported (Fei et al., 2016). For translation inhibition experiments, cells were incubated with puromycin (100ug/mL) for 30 minutes prior to AHA treatment. Nutlin3A treatment was for 36 hours at the indicated doses. For Caspase-3 experiments, cells were treated with Actinomcycin D (5ug/mL) for 12 hours. For siRNA experiments, cells were treated 25pmol smartpool siRNA (Dharmacon) and Lipofectamine RNAiMax transfection reagent (Thermo). Knock-down efficiency was assessed after 6 days by western blotting. The following siRNA pools were used: NPM1 (M-015737-01-0005), IPO7 (L-012255-00-0005), TP53 (L-003329-00-0005), RBM10 (E-009065-00-0005). Hoxb8-immortalized myeloid cells were generated as previously described (Wang et al., 2006) and cultured in RPMI 1640 supplemented with 10% Fetal bovine serum (FBS), 1% murine stem cell factor (SCF) and 1µM β-Estradiol. The cells were grown in 1µg/mL doxycycline to induce the U2AF1 transgene expression. Cells were induced to differentiate by culturing in the absence of β-Estradiol for 6 days before cells were stained for CD11b (Biolegend) and analyzed on the LSRII flow cytometer (BD Biosciences). Data analysis was performed using FCS Express (De Novo Software).

### Polysome profile analysis

The following steps are used to compute changes in translation efficiency from polysome sequencing profiles measured in Palangat et al., 2019:

1. For each sample (wt/wt, wt/s34f, wt/s34f-), 12 fractions were collected from the sucrose gradient, and the bottom/heaviest 10 fractions were sequenced.
2. Fractions 10-12 were pooled to increase coverage, resulting in 8 fractions in the final analysis. Fraction 5 and 6 correspond to the monosome fractions.
3. Reads mapping to particular message are normalized by the total reads in that sample and multiplied by 1×10^6^, generating counts per million (CPM) for each transcript in each fraction.
4. The CPM in each fraction constitutes the polysome profile (poly(x), where ‘x’ designates the sucrose fraction) which is now normalized for read depth across fractions/samples.
5. This profile can be further normalized by a weighting function which accounts for the non-linear relationship between sedimentation coefficient, position in the sucrose gradient, and molecular mass: S ∼ x^3/2^, where *S* is the sedimentation coefficient in Svedrup (Marks, 2001). Here, we use the following weights for each fraction after pooling: *w*=[0.6, 1.0, 1.6, 2.2, 2.9, 3.7, 4.5, 19.0].
6. The final numeric value, reflecting translation efficiency, is Σw_i_poly(x_i_). This number, generated for each transcript in each of the three cell lines, is the basis for clustering in Fig. 1A. Hierarchial clustering was done using ClustVis (Metsalu and Vilo, 2015).

### Gene Set Enrichment Analysis

GSEA analysis was done with GSEA v. 4.0.3 from The Broad using the ‘GSEA on a pre-ranked gene list’ option. Gene set database is c5.bp.v7.0.symbols.gmt run with 1,000 perturbations. The enrichment statistic is ‘weighted.’ Accepted gene sets contained between 15 and 500 members.

### Luciferase assay

Luciferase plasmids were introduced by lentiviral transduction. Equal number of cells were plated in a 96 well dish. After 24 hours, cells were treated with the Dual-Luciferase Reporter Assay System as per manufacturer’s instructions (Promega). Luciferase activity was measured by Victor plate reader (Perkin Elmer).

### AHA metabolic labeling

Global translation was assessed by AHA pulse labeling as previously described (Tom Dieck et al., 2012) with the following modification. Cells were plated on coverslips (VWR). After 24 hours, cells were pulsed with AHA for 1 hour. Cells were then fixed in 4% paraformaldehyde in PBS, permeabilized in 0.1% TritonX in PBS and blocked in blocking reagent that contains 2% BSA and 5% donkey serum in PBS for 3 hours. Cells were washed once in PBS. The click chemistry reaction was composed of Tris(2-carboxyethyl)phosphine (TCEP) (Thermo), Tris[(1-benzyl-1H-1,2,3-triazol-4-yl)methyl]amine (TBTA) (Sigma). The reaction was allowed to proceed for 1 hour. Cover slips were washed in 0.1% TritonX in PBS 3 time, 15 minutes each and mounted microscopy slides using DAPI-containing mounting solution.

### Quantitative Mass Spectrometry

Cells were lysed and total protein was quantified using BCA reagent (Thermo). The samples were reduced using TCEP (Thermo), alkylated with iodoacetamide (Sigma) and trypsinized (Promega) overnight. TMT tags were attached to the peptides and ran on MS (Thermo).

### Image acquisition and analysis

Slides were imaged using a custom epifluorescence microscope equipped with a Plan-Apochromatic 40X (NA 1.4) objective (Zeiss), illuminated with a LED light source (Model Spectra-6LCR-SA, Lumencor) and the emitted fluorescence collected with a CMOS camera (Hamamatsu ORCA-Flash 4.0). The microscope was controlled using µManager (Edelstein et al., 2014). A custom pipeline was employed to compute total fluorescence intensity per cell using Cellprofiler (Broad Institute) (Carpenter et al., 2006).

### Western Blotting

Cells were lysed in 1x RIPA buffer, boiled in SDS and run on SDS-PAGE. Proteins were transferred onto nitrocellulose membrane, blocked in 5% milk and probed with the following antibodies: NPM1 (Abcam, 1:1000), IPO7 (Abcam, 1:1000), TP53 (Bethyl, 1:500), Cyclin A2 (BD Biosciences, 1:1000), Tubulin (Abcam, 1:1000), Actin (Sigma, 1:10000).

### Viability assay

To assess changes in cell viability, cells were counted using an automated cell counter (BioRad) and equal number of cells were plated in a 96 well plate. On the sixth day of the siRNA experiment, fresh medium was added that contained 1:10 dilution of the WST1 reagent (Sigma). Cells were incubated for different time points before the absorbance was measured at 450 nm using a plate reader (BioRad).

### Apoptosis assay

Apoptotic induction was examined by checking for Active Caspase-3 levels using a PE-conjugated anti-Caspase-3 antibody per the manufacturer’s instructions (BD Biosciences). PE levels were examined using an LSRII flow cytometer (BD Biosciences). Data analysis was performed using FCS Express (De Novo Software).

### Cell cycle analysis

Cell cycle progression was examined using the Click-it Alexa Fluor 647 kit (Thermo). Briefly, cells were plated on cover slips. After 24 hours, cells were pulsed in Edu for 10 minutes and then fixed in 4% paraformaldehyde in PBS, permeabilized in 0.1% TritonX in PBS. Cells were then washed twice in 2% BSA in PBS. The EdU was reacted with azide-conjugated fluorophore in the presence of Copper sulfate. The click chemistry reaction was allowed to proceed for 1 hour. Cells were washed twice in 2% BSA in PBS. DNA was labelled using Hoechst at a dilution of 1:10000 mounted on Prolong Gold mounting solution (Thermo). The fraction of cells in each stage of the cell cycle was quantified using a custom Matlab script (Mathworks).

### Northern blotting

Equal numbers of cells were harvested for all conditions using the Luna Automated Cell Counter (Logos Biosystems). Total RNA was extracted using Trizol (Thermo) according to the manufacturer’s instructions. The RNA was separated on a denaturing gel and transferred to nylon membrane (GE healthcare) overnight as previously described (Akef et al., 2015). The membrane was hybridized to a ^32^P-labeled probe overnight and imaged using a Typhoon phosphoimager (Molecular Dynamics). To probe for the ITS2, an oligonucleotide was 5’-labeled using PNK enzyme (NEB). The ITS2 probe was previously reported (Donati et al., 2013) and its sequence is GAGGGAGGAACCCGGACCGCAGGCGGCGGCCACGGGAACTCGGCCCGAGCCGGCTCTCTC. The 5’ ETS probe sequence was previously reported (Fumagalli et al., 2009) and its sequence is GGCGAGCGACCGGCAAGGCGGAGGTCGACCCACGCCACACGTCGCACGAACGCCTGTC. For GAPDH probe, the pET30-2-GAPDH plasmids was a gift from David Sabatini (Addgene plasmid # 83910) (Pacold et al., 2016) and gene-body labeling was performed using the Prime-A-Gene kit (Promega).

### OPP labeling

Immortalized myeloid cells were treated with 10µM OPP (Click Chemistry tools) for 1 hour. Cells were fixed and permeabilized. Click chemistry was employed to conjugate a fluorophore to the OPP containing polypeptides according to the manufacturer’s instructions (Thermo). Cells were washed in PBS and analyzed on the LSRII flow cytometer (BD Biosciences). Data analysis was performed using FCS Express (De Novo Software).

### Mutual Exclusivity of U2AF1 and NPM1 mutations

Cancer patient data was retrieved from cBioPortal (www.cbioportal.org) as of October, 4^th^, 2019. All adult AML and MDS patient studies were selected. These include: “Acute myeloid leukemia or myelodysplastic syndromes (Wash U, 2016)”, “Acute myeloid leukemia (OHSU, 2018)”, “Acute myeloid leukemia (TCGA, PanCancer Atlas)”, and “Myelodysplasia (UTokyo, Nature 2011)”.

## Acknowledgements

This work was supported by the intramural research program of the National Cancer Institute. Thorkell Andresson and Sudipto Das of the NCI Mass Spectrometry Institute carried out the MS data collection and initial analysis. We thank Dr. Jeff Coller for sharing rRNA northern blotting protocols. We also thank members of the Larson lab for comments on the manuscript. The authors declare no competing financial interests.

## Author contributions

AA and DRL conceived and designed the study. AA performed most experiments and analyzed data. DRL analyzed the polysome profiling data. SDC contributed software and advice for cell cycle analysis. KM contributed reagents. AA and DRL wrote the manuscript with feedback from SDC.

## Supplementary Figure Legends

**Figure S1.**
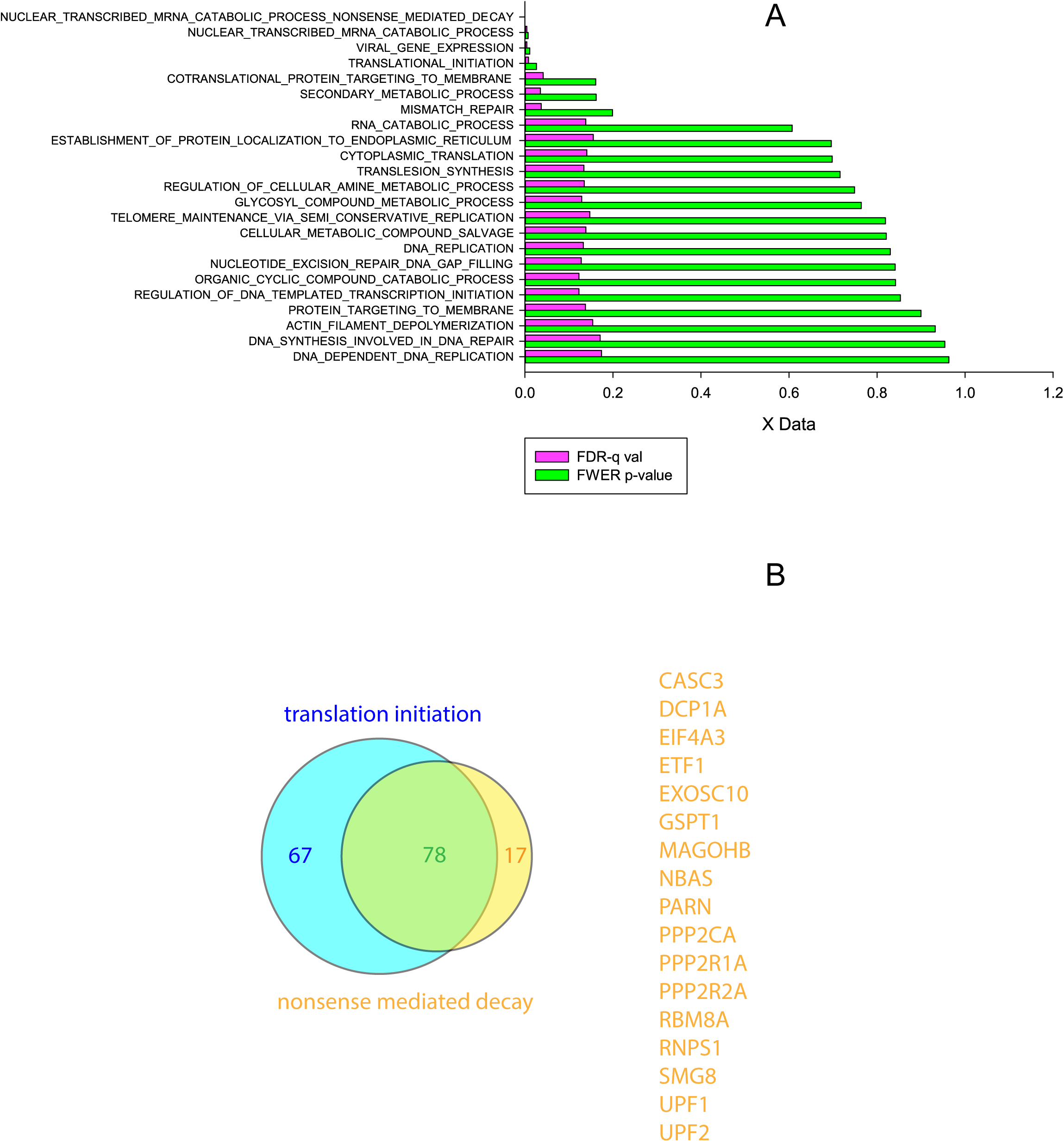
Full results of the GSEA analysis on MS data. A) The top four GO categories show similar FDR and FWER and contain many of the same genes. B) The overlap between the GO categories of “nuclear transcribed mRNA catabolic process nonsense mediated decay” and “translation initiation” is shown as a Venn diagram. The 17 genes which are present in “nuclear transcribed mRNA catabolic process nonsense mediated decay” but not “translation initiation” are listed.

**Figure S2.**
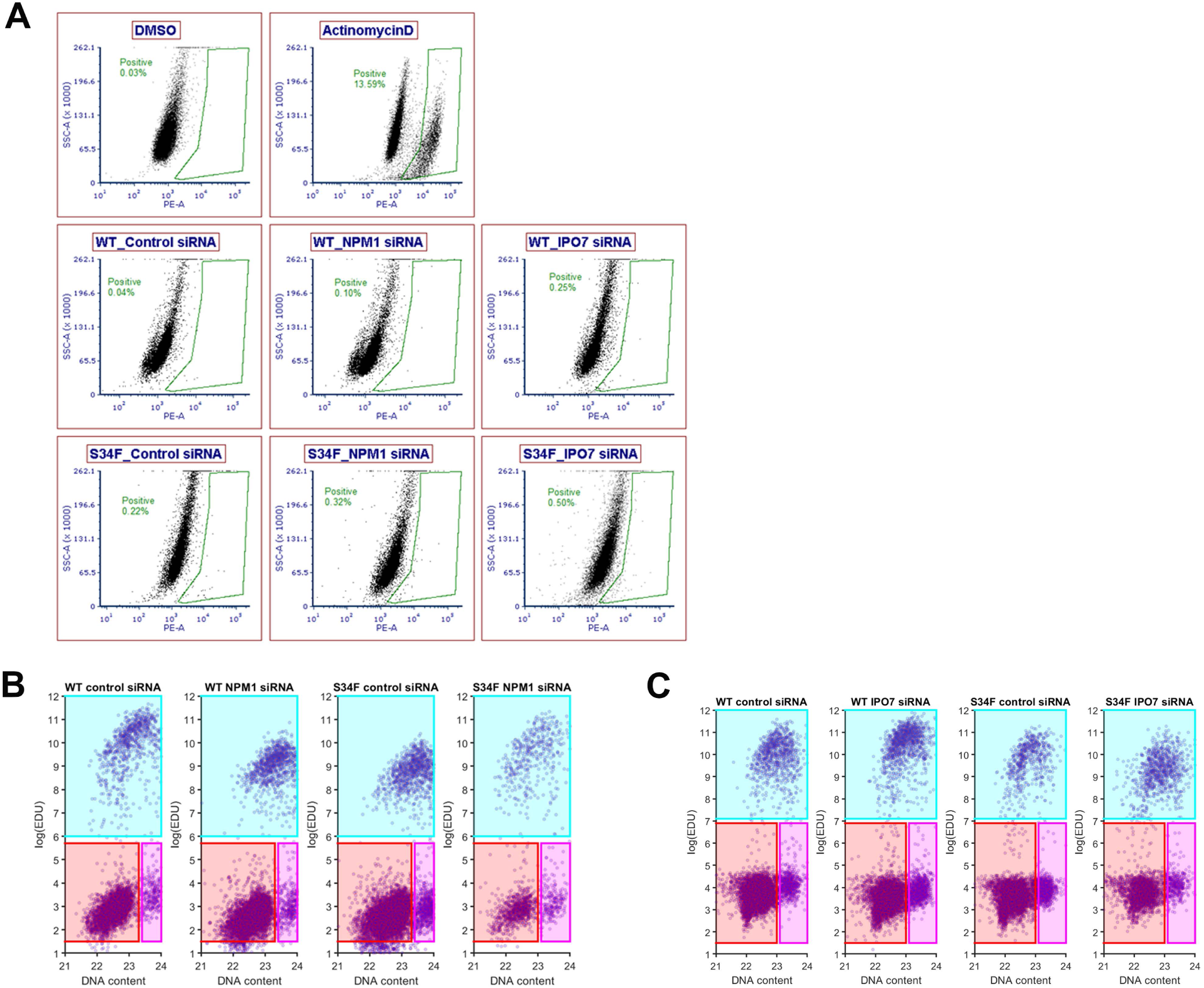
Gating Strategy from representative Caspase-3 and EdU labeling experiments.

